# Cross-task implications: How hippocampal event boundary responses predict unrelated memory performance

**DOI:** 10.1101/2024.10.02.616238

**Authors:** Daphne van Dijk, Silvy H.P. Collin

**Affiliations:** Tilburg School of Humanities and Digital Sciences, Tilburg University Tilburg, Netherlands

**Keywords:** event segmentation, hippocampus, machine learning

## Abstract

Hippocampal responses at event boundaries have been shown to predict memory performance for these events. However, are these hippocampal event boundary responses specific to memory for those particular events, or can they also have predictive power across various memory tasks? We used data from the Cam-CAN project (fMRI data from continuous movie viewing and memory results from an unrelated Famous Faces Task, N = 630) to determine whether hippocampal responses at event boundaries during the continuous movie viewing were indicative of memory performance in the unrelated Famous Faces task using various machine learning algorithms. The results showed that memory performance in the Famous Faces Task could be predicted based on participants’ hippocampal event boundary responses in another task, which suggests that the hippocampal event boundary responses are indicative for general memory performance. This might indicate importance of these hippocampal event boundary responses in terms of general information processing of the human brain.

## 1 Introduction

When people experience everyday activities, they parse this stream of activity into discrete, meaningful events. According to Kurby and Zacks (2008), an ‘event’ can be described as a segment of time at a certain location where you can indicate a clear beginning and end. The understanding and perception of events is supported by so-called ‘event models’ that make predictions about what will happen next. When something happens that is not consistent with the prediction based on the current event model, it will be experienced as a prediction error. Subsequently, the current model gets an update based on the latest perceptual information. The time-points at which these updates are made are referred to as ‘event boundaries’ (Kurby & Zacks, 2008). When people are explicitly asked to mark event boundaries in, for example, a movie, it appears that people strongly agree on the location of these event boundaries (Newtson, 1973). This phenomenon was also explored in fMRI-studies. For instance, in Zacks et al., 2001, people were shown tapes of everyday events in the MRI-scanner. Participants were unaware of the segmentation task during this stage. When these same people were later explicitly asked to mark event boundaries in these same videos, there appeared to be a significant correlation between transient changes in brain activity and the explicitly self-labeled event boundaries. These results suggest that event segmentation is a natural and spontaneous aspect of human information processing (see also Zacks and Swallow, 2007 and Geerligs et al., 2021). Furthermore, increased hippocampal activity is specific and sensitive in its response to subjective event boundaries. Moreover, this activity was larger at those boundaries for which they found high consensus across participants (Ben-Yakov & Henson, 2018). This means that strong, obvious event boundaries also trigger stronger responses in the hippocampus. In this study, they recorded activity peaks in the hippocampus and then analyzed the alignment between the event boundaries and these activity peaks. These event boundaries were identified by an independent group of participants. This analysis demonstrated that increased hippocampal activity was highly correlated with the identified event boundaries (Ben-Yakov & Henson, 2018). In summary, these studies have shown that brain activity seems to be modulated by event structures and that there is general agreement on those structures among people (Ben-Yakov & Henson, 2018; Newtson, 1973; Zacks et al., 2001). Moreover, it has become clear that event boundaries are reflected by peaks in hippocampal activity (Barnett et al., 2024; Ben-Yakov & Henson, 2018; Bilkey & Jensen, 2021; Brunec et al., 2018; Griffiths & Fuentemilla, 2020; Reagh et al., 2020; Zacks et al., 2001).

Numerous previous studies indicated that event segmentation of ongoing activity plays an important role in people’s ability to remember and understand things (Aitken & Kok, 2022; Baldassano et al., 2017; Bein et al., 2020, 2021, 2023; Ben-Yakov & Henson, 2018; Buckley et al., 2022; Geerligs et al., 2021; Güler et al., 2024; Kalbe & Schwabe, 2020; Kurby & Zacks, 2018; Nolden et al., 2024; Pettijohn & Radvansky, 2016; Pradhan & Kumar, 2022; Radvansky & Zacks, 2017; Sargent et al., 2013; Sinclair et al., 2021). Baldassano et al. (2017) discovered a relationship between event boundaries and hippocampal encoding in a movie-viewing experiment. Their research showed that the hippocampus was triggered at the end of an event, i.e., at event boundaries, to encode this new information about this event into memory. This implies that the segmentation of events contributes to how memories are organized in memory. The encoding seemed to be most powerful when the activity in the hippocampus was relatively low during an event, but considerably high at an event boundary (Baldassano et al., 2017). Kurby and Zacks (2018) also note that the event segmentation process is used to update human working memory and regulate encoding in people’s long-term memory. Worse event memory can therefore be explained by bad event segmentation ability. Event segmentation ability can be defined as the level at which someone agrees with a larger sample regarding the location of the event boundaries in ongoing activities (Sargent et al., 2013). According to Kurby and Zacks (2018), if people cannot segment events properly, these events will not be properly encoded and this in turn has a negative effect on memory recall. Sargent et al. (2013) showed that the ability to segment a continuous experience can accurately predict memory related to this specific activity. In this study, people who were better at segmenting a movie were able to remember more actions from this movie afterwards. In conclusion, the ability to properly segment ongoing activity is crucial for memory, but all of these studies focus on event-specific memory of that particular task. It is less clear whether a reduced event segmentation ability, which would be reflected by less clear hippocampal peaks at event boundaries, is also indicative of people’s memory performance on a more general level.

Thus, it has been shown multiple times that there is a coincidence of activity peaks in the hippocampus and event boundaries, however, less is known about whether these hippocampal peaks at event boundaries can also be a reliable “more general” predictor across-tasks. The purpose of this study is therefore to extend the findings of previous studies by gaining more insight into the interaction between event segmentation, hippocampal activity, and the performance on a general and unrelated memory test. This will be accomplished by applying machine learning techniques (i.e., linear regression, support vector machine, multilayer perceptron) as well as intersubject correlation analyses (ISC, Hasson et al., 2004). Our goal is to determine whether hippocampal time courses on one cognitive task can be used to make reliable predictions for people’s memory performance across different tasks. If the hippocampal responses to event boundaries appear to be a reliable predictor of the performance on another unrelated task, it would suggest that a reduced event segmentation ability is not only indicative of someone’s event-specific memory, but also of someone’s memory performance in general.

Altogether, we hypothesize that participants that have hippocampal peaks that are better aligned with event boundaries have better overall information processing, and should outperform also in other information processing tasks. Thus, with this study we answer the following research question: *To what extent are hippocampal responses to event boundaries in an ongoing activity indicative of across-task memory performance?*

## 2 Methods

### 2.1 Participants

Data used for this project was obtained from the Cam-CAN repository (available at http://www.mrc-cbu.cam.ac.uk/datasets/camcan/) from the Cambridge Center for Aging and Neuroscience (University of Cambridge 2010; Shafto et al., 2014; Taylor et al., 2017). Cam-CAN uses cognitive, neuroimaging and epidemiological data to learn more about how elderly can best maintain cognitive capacities. The following material was used from this dataset for the current study: the fMRI data from the movie-viewing experiment (N = 649), and the behavioral data from the Famous Faces task (N = 660). In total there were 631 participants who took part in both movie-viewing as well as the Famous Faces task. One participant was excluded because the behavioral data for this participant was incomplete. Thus, the final dataset consisted of 630 participants. Among the participants were 315 men and 315 women with a mean age of 54.88 (SD = 18.31, range from 18.5 to 88.9 years old).

### 2.2 Task

#### 2.2.1 Movie-viewing task

Participants who took part in this movie-viewing task watched a condensed 8-minute version of Alfred Hitchcock’s “Bang! You’re Dead” in the fMRI-scanner. Even though the full 25-minute video was considerably shortened, the narrative of the video was preserved.

#### 2.2.2 Famous Faces task

The Famous Faces task has been selected to serve as the measure of general memory performance (i.e., unrelated to the movie-viewing task). This semantic memory test measures participants’ ability to recognize famous people from photos. Participants were shown photos of 30 celebrities and 10 random unknown people. Participants were asked if they recognized the person in the photo. If so, they were asked if they knew this person’s name, and if they could provide occupational information about this person.

### 2.3 Description of fMRI data used for this study

#### 2.3.1 Movie-viewing task

We used the pre-processed fMRI-data of the movie-viewing task for this study. Details about the fMRI data in terms of data collection and pre-processing can therefore be found in the paper written by Taylor et al. (2017). In short, the fMRI data were collected using a 3T Siemens TIM Trio equipped with a 32-channel head coil. T1-weighted images as well as T2*-weighted echo planar images (EPIs) were obtained. A multi-echo sequence (whole brain coverage; TR = 2470 ms) was used. The acquired functional and structural images were pre-processed using SPM12 (see http://www.fil.ion.ucl.ac.uk/spm) as implemented in the Automatic Analysis pipeline system described in the paper by Cusack et al. (2015). In short, the obtained functional images were corrected for slice-timing differences and motion. An anatomical group template was created using the DARTEL procedure and then transformed into Montreal Neurological Institute (MNI) space. The EPI images were co-registered to the T1 image and normalized into MNI space.

### 2.4 Data analysis

#### 2.4.1 Famous Faces task

We approached the classification in this study with binary classifiers (“good” vs “bad” memory performance) on the Famous Faces task.

Shafto et al. (2014), made a distinction between four memory components: (1) number of famous faces recognized, (2) number of faces for which occupational information was given, (3) number of faces whose full names were given, and (4) correct rejections (i.e., the number of unfamiliar faces that were correctly identified as unfamiliar). We used component 2 and 3 separately as memory scores, as well as a memory score for which we subtracted the percentage of false alarms (calculated based on memory component 4) from the percentage correct recognition (component 1). Additionally, the “final score” used is the sum of these three scores (descriptive statistics in table 1).

**Table 1:** Descriptive statistics per memory component.

| component | min | max | mean | SD | median |
| --- | --- | --- | --- | --- | --- |
| true recognition | 20 | 100 | 87.56 | 14.64 | 90 |
| naming | 4 | 30 | 23.58 | 5.79 | 25 |
| occupation | 6 | 30 | 26.86 | 4.20 | 28 |
| final score | 39 | 160 | 138.00 | 21.79 | 144.67 |

To separate participants in “good” and “bad” memory performance, we used the median. The distribution of the participants over the two classes per memory aspect is shown in figure 1.

**Fig. 1.**
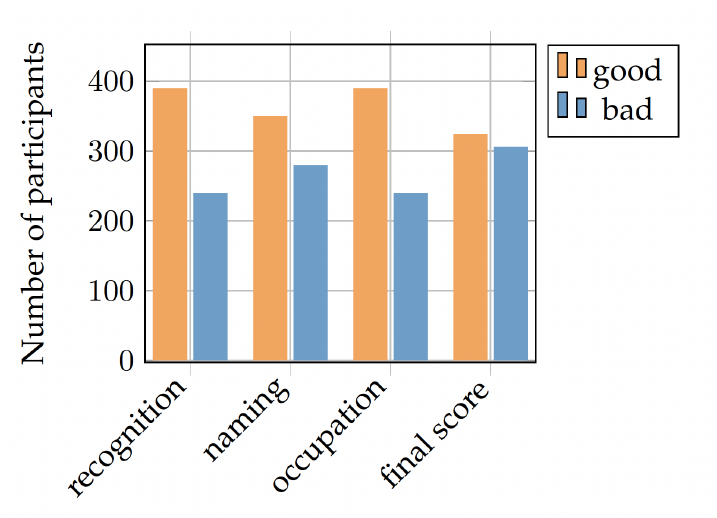
The distribution across “good” and “bad” memory performance for each of the 4 memory scores used in the study

### 2.4.2 Machine learning analyses

#### Algorithms and evaluation metrics

We used three algorithms in this study: logistic regression, support vector machine, and multilayer perceptron. The algorithms were implemented and defined using the scikit-learn library (version 0.23.2). Several kind of evaluation metrics were implemented to evaluate and compare the classification performance of the different machine learning models. The models were trained using accuracy, but other metrics are preferred for evaluating classification performance with unbalanced sample sizes (Arbabshirani et al., 2017). The get a thorough overview of model performance we also calculated precision, recall, F1-score and the Area Under the Receiver Operating Characteristic Curve (ROC AUC). Chance level is 50 percent (“good” vs “bad”) on all algorithms.

#### Hippocampal timecourses

In this study, we only used the hippocampus (of the left and right hemisphere). Since no difference was expected between the left and right hemisphere, the average of these two sequences was used to represent people’s hippocampal time course. The obtained hippocampal time courses were normalized (z-scored) within subjects to enable better comparison across participants.

#### Cross-validation

The z-scored mean array of the activity in the hippocampus of the left and right hemisphere was used as input in three different algorithms. For these three algorithms, it was examined whether they can be used to correctly predict four components of people’s performance on the Famous Faces task. The dataset was divided into 80 percent training data and 20 percent test data in a stratified fashion. The models were trained using k-fold cross-validation (k = 5) to make optimal use of the available data.

#### Hyperparameter tuning

To find the optimal hyperparameters, a grid search was performed per algorithm during the training on the binarized final memory score. The selected hyperparameters based on these grid searches were then applied to all models for the individual memory components. As a result, the same hyperparameters were used for each memory component per algorithm. Thus, the hyperparameters only vary between algorithms and not across memory components. This makes it possible to interpret the classification performance on the various memory components more unambiguously. The hyperparameters that gave the best results can be found in Table 2.

**Table 2:** The selected hyperparameters per algorithm based on performed grid searches. For the hyperparameters not mentioned, the default settings of the scikit-learn library (version 0.23.2) have been applied.

| Algorithm | hyperparameters |
| --- | --- |
| LR | 'C': 1, 'penalty': 'l1', 'solver': 'saga' |
| SVM | 'C': 1, 'cache_size': 100, 'kernel': 'rbf' |
| MLP | 'activation': 'logistic', 'hidden_layer_sizes': 1000, 'solver': 'adam', 'alpha': '0.001', 'learning_rate': 'adaptive' |

#### Comparing across machine learning models

Furthermore, to test whether there were significant differences between the algorithms in their classification performance, Cochran’s Q Tests were performed for each memory component using the accuracy scores achieved on the test dataset. Cochran’s Q test is a non-parametric statistical technique to compare the performance of multiple machine learning models (Raschka, 2020). It tests the null hypothesis that there is no significant difference between multiple classification models in their accuracy on the same test set.

#### 2.4.3 Inter-subject correlation analysis

As an additional test, it was examined whether there was a significant difference between the level of event segmentation consistency between the good-memory group and the bad-memory group. This was established by performing an intersubject correlation (ISC) analysis on the hippocampal time courses (complete dataset). Intersubject correlation (ISC) analysis of functional brain imaging data offers great insight into how brain activity is correlated across different participants when they are exposed to the same ongoing activity (e.g., movie-watching). This means that ISC quantifies the consistency of neural responses to these kind of naturalistic stimuli among people (Hasson et al., 2004; Nastase et al., 2019). The ISCs can be established by computing the correlation coefficients between all possible pairs of participants. According to Nastase et al. (2019), statistical inference for ISC analysis is complicated as each participant contributes to the calculation of the ISC of all other participants. For that reason, the assumption of independence is violated and standard parametric tests (e.g. T-tests) are not suitable for this type of analysis. Therefore, subject-wise permutation tests are used for comparing two groups that are expected to have different ISC values (Chen et al., 2016). Statistical significance is determined by testing against a null distribution resulting in a reliable permutation-based p-value corresponding to the two-sided test (Chen et al., 2016; Nastase et al., 2019). Here, the ISCs were computed using a pairwise approach. Next, a two-sample Monte Carlo approximate permutation test was performed on these ISCs using 1000 iterations. The labels belonging to the binarized final memory score were used as the group labels. In this way, differences were computed between the median ISC-score for within-group correlations while the between-group correlations were excluded. Monte Carlo resampling had to be applied because an exact test would result in an infinitely long list of possible permutations. After all, in a two-sample test the number of possible permutations equals the factorial of N (N = 630 in this case).

#### 2.5 Software

For this study, all (pre)processing steps were executed by using Python (version 3.7.3) in Jupyter Notebooks (version 6.1.1). The MATLAB-files from the received Cam-CAN dataset were converted to Python format using the SciPy library (version 1.5.2). The algorithms were implemented and defined using the scikit-learn library (version 0.23.2). Furthermore, NumPy (version 1.19.1), pandas (version 1.1.1), glob2 (version 0.7) and Matplotlib (version 3.3.1) were used for data preprocessing, data visualization, and the implementation of the algorithms. The Cochran’s Q Tests were implemented using the Mlxtend library for Python (version 0.17.3). The intersubject correlation analysis was performed using the Brain Imaging Analysis Kit for Python (see http://brainiak.org). The implementation of this analysis kit is based on the ISC tutorial of Nastase (2019).

## 3 Results

### 3.1 Classification

We used the hippocampal activity at event boundaries to predict memory performance at the Famous Faces task (i.e., a memory task unrelated to the neural data) using linear regression, support vector machine and multilayer perceptron. For all three algorithms, we ran a separate model for each of the 4 determined memory scores (see Table 3).

**Table 3:** The classification performance on the test dataset per algorithm for the different memory components. Test performance was evaluated based on accuracy, recall, precision, F1-score and the ROC AUC score.

| component | models | accuracy | recall | precision | F1-score | ROC |
| --- | --- | --- | --- | --- | --- | --- |
| recognition | LR | 0.57 | 0.46 | **0.37** | **0.41** | 0.57 |
|  | SVM | **0.64** | **0.73** | 0.16 | 0.26 | **0.62** |
|  | MLP | 0.61 | 0.60 | 0.12 | 0.20 | 0.61 |
| naming | LR | 0.59 | 0.51 | **0.51** | **0.51** | 0.59 |
|  | SVM | **0.60** | 0.54 | 0.38 | 0.44 | **0.60** |
|  | MLP | **0.60** | **0.57** | 0.23 | 0.32 | 0.57 |
| occupation | LR | 0.62 | 0.50 | **0.46** | **0.48** | 0.58 |
|  | SVM | **0.63** | **0.60** | 0.12 | 0.21 | **0.62** |
|  | MLP | 0.56 | 0.40 | 0.29 | 0.34 | 0.58 |
| final score | LR | 0.53 | 0.53 | **0.51** | 0.52 | 0.51 |
|  | SVM | **0.62** | **0.66** | 0.49 | **0.56** | **0.66** |
|  | MLP | 0.56 | 0.59 | 0.35 | 0.44 | 0.60 |

All models scored above 50 percent chance level indicating that we could indeed predict memory performance of the Famous Faces task using hippocampal responses of an unrelated movie-viewing task. The SVM model seemed to perform best in most cases (between 0.60 and 0.64 for all 4 memory components). Also when investigating the recall scores for all 4 memory components, it was especially the SVM that managed to predict memory performance based on the hippocampal data (especially for the recognition performance with 0.73 and the final score with 0.66). However, when only precision scores were considered, the SVM and MLP appeared to perform very poorly. Thus, the LR model turned out to be the best classifier in terms of precision scores. No major differences were observed between the different memory components.

### 3.2 Comparing across models

We used Cochran’s Q Tests for each memory component using the accuracy on the test dataset to determine if there were significant differences between the three algorithms in terms of classification accuracy. No significant difference was found between the classifiers for any memory component (see Table 4).

**Table 4:** Results of Cochran’s Q Tests: the Q-value and associated p-value per memory component.

| component | Q-value | p-value |
| --- | --- | --- |
| recognition | 2.450 | 0.295 |
| naming | 0.195 | 0.907 |
| occupation | 2.577 | 0.276 |
| final score | 4.217 | 0.121 |

### 3.3 Intersubject correlation (ISC) analysis of hippocampal time courses

The intersubject correlation analysis revealed a median correlation of 0.439 across all values. A two-sample Monte Carlo approximate permutation test was conducted to compare the ISCs in the bad-memory group with the good-memory group. The actual observed group difference in terms of the median ISC values turned out to be −0.088. The non-parametric test showed that there was a significant difference (p = 0.014) in the median ISC values for the bad-memory group (0.393) and the good-memory group (0.480). Thus, the time courses correlated significantly stronger in the good-memory group compared to the bad-memory group. This also means that there is more consistency in the time courses in the good-memory group than in the bad-memory group.

## 4 Discussion

The purpose of this current study was to find out whether hippocampal responses to event boundaries are indicative of general, across-task memory performance. This was examined by comparing three different algorithms in their ability to predict people’s performance on the Famous Faces task using hippocampal time courses of continuous movie-viewing. As an additional test, an intersubject correlation (ISC) analysis was carried out to gain insight into the group differences in the hippocampal time courses between people with good versus bad memory performance. All models used (LR, SVM, MLP) were able to predict performance on the unrelated (Famous Faces) task based on the hippocampal timecourses drawn from the movie viewing task.

### 4.1 Differences across machine learning models

We also thoroughly investigated possible differences across the three algorithms used. In general, it can be concluded that there were no significant differences across the three algorithms in their prediction ability in this study. When looking at accuracy, SVM seemed to perform slightly better than the other two models. However, completely different conclusions could be drawn by only looking at the precision score and the F1-score. The precision scores and F1-scores for the SVM models were in fact considerably low compared to the scores on the other evaluation metrics. The MLP models also suffered from this issue, but the LR models did not. It does make sense that this problem arises. After all, only healthy people took part in this study. This means that there were probably relatively few good representative examples in the dataset of people with really poor memory capacity. It has been decided to set the threshold at the median final score to minimize class imbalance, but the disadvantage of this approach is that some people labeled with “bad” memory may in fact have quite normal / average memory performance. As a consequence, the hippocampal time course of these people will presumably not deviate much from many people in the good-memory group. This probably made it more difficult for algorithms to learn the relationship between the “bad” memory label and the hippocampal time courses. This issue may also explain why the SVM and MLP model show a large difference between the precision scores and the scores on the other evaluation metrics. After all, these models are nonlinear and probably created complex decision boundaries, with many “average” observations ending up in the bad-group because those were almost identical to the observations that truly belonged to the bad-group. The LR model uses a less complex decision boundary since it is a linear classifier. With the LR model, observations were classified almost at chance-level. The difficulty the model has in finding the right location for a linear decision boundary could possibly explain this. In this situation, some observations will incorrectly end up in the good-group and some incorrectly in the bad-group. While in a nonlinear model there will be a tendency to classify too many observations as “bad”, and this will result in poor precision scores. However, this explanation is speculative and must be further investigated in follow-up research.

### 4.2 Predictive power of hippocampal timecourses

All these results together make it plausible that the hippocampal responses to event boundaries are related to general memory capacity. First of all, algorithms were capable of classifying memory performance using hippocampal time courses better than one would expect based on chance alone. Secondly, there appeared to be a significant difference in event segmentation consistency between the two groups. Thus, what someone’s hippocampal time course of ongoing activity looks like provides information about this person’s overall memory capacity. However, given the research approach in this study, it can never be established with certainty that it is actually only the event boundaries that are indicative of the memory performance. The indicative factors in the time course could involve other things as well. However, since Ben-Yakov and Henson (2018) indicated that activity peaks in this exact same dataset are specific to event boundaries, it is extremely likely that the event boundaries are indeed indicative factors. Despite the significant difference in hippocampal activity between the two groups, classification performance turned out to be only slightly better than the baseline. A weak classification rate like this is not uncommon in this type of research. According to Arbabshirani et al. (2017), this can be explained by the fact that the time courses of the different groups usually overlap to a great extent. As a consequence, a significant group difference does not necessarily guarantee a strong classification performance.

What is also striking in these results is that the MLP model only yields the best results in a very few cases. In previous studies, the use of neural networks has proven to be very effective for neuroimaging classification (e.g., Güçlü and van Gerven, 2017; Wen et al., 2018). However, most deep learning studies have used more complex neural network architectures such as Convolutional Neural Networks (ConvNet / CNN) or Recurrent Neural Networks (RNN). The MLP is one of the most basic versions of a neural network. It is possible that the classification of complex fMRI data may require these type of more complex neural networks. In addition, neural networks perform especially well when a large amount of training data is available (Sun et al., 2017; Ulloa et al., 2018). It is therefore likely that the relatively weak results of the MLP model are (partly) caused by the relatively small size of the dataset.

In summary, this study suggests that hippocampal peak activity at event boundaries in ongoing activity relate to general across-task memory performance. This extends findings in earlier work of hippocampal peak activity being related to task-specific memory performance (Aitken & Kok, 2022; Baldassano et al., 2017; Barnett et al., 2024; Bein et al., 2020, 2021, 2023; Ben-Yakov & Henson, 2018; Bilkey & Jensen, 2021; Brunec et al., 2018; Buckley et al., 2022; Geerligs et al., 2021; Griffiths & Fuentemilla, 2020; Güler et al., 2024; Kalbe & Schwabe, 2020; Kurby & Zacks, 2018; Nolden et al., 2024; Pettijohn & Radvansky, 2016; Pradhan & Kumar, 2022; Radvansky & Zacks, 2017; Reagh et al., 2020; Sargent et al., 2013; Sinclair et al., 2021). This might reflect that someone’s ability to detect event boundaries is reflective of a general benefit in information processing skills. However, additional research is needed to be able to make more solid statements about it.

### 4.3 Future research

For future studies, it would be good to make use of larger, strictly demarcated groups. By also collecting data from people suffering from memory deficits, it is easier to investigate differences in the hippocampal time courses since this is expected to result in less overlap in the time courses. This will allow stronger conclusions to be drawn about the relationship between hippocampal activity and overall memory capacity. Moreover, the amount of data in this dataset is not sufficient to achieve great success with deep learning methods. More data could improve this, and it would also reduce the risk of overfitting. Furthermore, it could also be very interesting to experiment with more complex neural network architectures. A major problem here is that it is often difficult to acquire a large (labeled) medical imaging dataset due to the high costs (Sun et al., 2017). In follow-up research, it may therefore be interesting to explore ensemble learning strategies in which models are used with low precision performance but relatively higher recall performance. If these models are diverse enough, it most likely means that the false positives will be diverse as well. This then makes it possible to cancel those false positives out by averaging the models (Ma et al., 2021). Ma et al. (2021) showed that this approach can yield great results when dealing with small classes. It has also been shown in other research areas that combining different classification models, including deep learning methods, can yield very good results (e.g., Kahou et al., 2016; Sun et al., 2017). Lastly, for future research it could also be interesting not to use the entire time course as input. For example, it is possible to choose to only investigate brain activity at the event boundaries that have a high inter-subject agreement. This will make it possible to determine with more certainty that the responses to event boundaries are indicative of memory performance because only boundary-evoked responses are studied, meaning that there will be less random noise in the data.

## 5 Conclusion

The results of this study show that all proposed classifiers were capable of predicting memory performance based on hippocampal time courses better than one would expect based on chance alone. Accuracy scores were found to be around 60 percent. These are not exceptionally high scores, but this is expected to be partly caused by various limitations as discussed above. These results suggest that hippocampal responses to event boundaries are indeed indicative of general across-task memory performance. This might indicate predictive power of these hippocampal event boundary responses in terms of general information processing of the human brain.

## Acknowledgements

Data collection and sharing for this project was provided by the Cambridge Centre for Ageing and Neuroscience (Cam-CAN). Cam-CAN funding was provided by the UK Biotechnology and Biological Sciences Research Council (grant number BB/H008217/1), together with support from the UK Medical Research Council and University of Cambridge, UK.

